# The nasopharyngeal virome in adults with acute respiratory infection

**DOI:** 10.1101/2023.08.21.554191

**Authors:** N.T. Sandybayev, V.Yu. Beloussov, V.M. Strochkov, M.V. Solomadin, J. Granica, S. Yegorov

**Author notes:** **Corresponding authors’ contact emails:** NTS, SY.

## Abstract

**Background:** Respiratory infections are a leading cause of morbidity and mortality worldwide, with a significant proportion thought to have a viral aetiology. Traditional diagnostic approaches often rely on targeted assays for “common” respiratory pathogens, leaving a substantial fraction of infections unidentified. Here, we used metagenomic next generation sequencing (mNGS) to characterize the nasopharyngeal virome associated with acute respiratory infection (ARI), with and without a positive PCR test result for a panel of common respiratory viruses.

**Methods:** Nasopharyngeal swabs from symptomatic outpatients (n=49), of whom 32 tested positive by a multiplex viral PCR, and asymptomatic controls (n=4) were characterized by mNGS. The virome taxa were stratified into human, non-human eukaryotic host, and bacteriophage sub-groups. We used a phage host classification as a proxy to establish bacterial taxa present in the nasopharynx. We then compared the virome biodiversity and presence of pathogens with known respiratory effects across the participant sub-groups.

**Results:** The nasopharyngeal virome exhibited similar diversity across the PCR-positive and - negative subsets. Among the top ARI-associated human viruses were enterovirus (16.3%, human rhinovirus, HRV-A), roseolovirus (14.3%, human betaherpesvirus 7, HBV-7) and lymphocryptovirus (8.16%, Epstein-Barr virus, EBV). The top three ARI-associated phage hosts were *Streptococcus* spp (32.7%), *Pseudomonas aeruginosa* (24.5%) and *Burkholderia* spp. (20.4%). The virome of both asymptomatic and symptomatic subjects was also abundant in the *Staphylococcus* (60.4%) and *Propionibacterium (Cutibacterium) acnes* bacteriophages (90.6%). The PCR and mNGS results were relatively concordant for human rhinovirus (HRV), but not for other PCR panel targets, including human parainfluenza (HPIV), adenovirus (HAdV), bocavirus (BoV) and seasonal coronavirus (HCoV).

**Conclusions:** mNGS revealed a high diversity of pathogens that could be cause to respiratory symptomatology, either as a single infection or a co-infection between viral and bacterial species. The clinical significance of the mNGS versus multiplex PCR findings warrants further investigation.

## Introduction

Most acute respiratory infections (ARI) are thought to have viral aetiology, although bacterial ARI are also common^1–3^ . Clinical management of uncomplicated ARI is typically empiric, especially in the limited-resource settings of rural regions or developing countries ^4,5^. Rarely, a laboratory-based characterization of ARI is done using microbiologic or molecular assays targeting “common” respiratory pathogens. Even then, over a third of ARI can remain unidentified, and it is often unclear whether the pathogens identified by targeted assays are the true cause of clinical symptoms ^1,6,7^.

This diagnostic gap has led to an increased interest in understanding the broader respiratory virome, encompassing not only well-known pathogenic viruses but also less characterized viral communities that may play a role in health and disease ^8,9^. The nasopharyngeal virome, in particular, represents a complex and dynamic ecosystem. Studies have revealed a diverse array of viruses, including human, non-human eukaryotic host, and bacteriophage sub-groups. This complexity extends beyond the presence of known pathogenic viruses, encompassing commensal viruses and interactions with bacterial communities that may influence respiratory health ^8,10^. Metagenomic next-generation sequencing (mNGS) has emerged as a powerful tool to explore the respiratory virome, offering an unbiased approach to detect known and novel viruses ^8^. However, the concordance between mNGS and traditional methods like multiplex PCR remains an area of active investigation, with varying results reported for different viral targets ^6^. The characterization of the nasopharyngeal virome holds potential implications not only for timely ARI diagnosis but also for development and deployment of effective therapies against specific ARI pathogens. The presence of co-infections between viral and bacterial pathogens and the role of less common viral and bacterial species in ARI patients from diverse demographic and geographic strata are some of the areas that require a better understanding. In our earlier work, we surveyed adult ARI outpatients in Kazakhstan during a period of low COVID-19 transmission in Spring 2021^5^. Globally, one hallmark of this period was a complete absence of influenza virus and reduced respiratory syncitial virus (RSV) transmission, but a resurgence of human rhinovirus (HRV)^11,12^. Consistently, in our cohort, 66% of the ARI outpatients were positive by a multiplex PCR for HRV, human parainfluenza virus (HPIV), adenovirus (HAdV) and seasonal coronaviruses (CoV) OCR-43/HKU-1, while a third of ARI cases were negative on the PCR ^5^.

Here, we expanded on these earlier results and further characterized the nasopharyngeal virome associated with acute respiratory infection (ARI) using metagenomic next generation sequencing (mNGS). Our study is part of the national initiative to enhance the clinical diagnostic capacity for ARI in the post-pandemic era.

## Materials and methods

### Study setting

This study is a follow-up to our earlier work exploring virologic causes of ARI among outpatients of two public hospitals in Kazakhstan ^5^. In this earlier study, 50 participants with ARI symptoms, who tested negative for SARS-CoV-2 and influenza A virus, were recruited in between May 18 and June 7, 2021. Written consent to participate was obtained from all participants and witnessed by the study coordinator. ARI was defined by the presence of at least one of the following respiratory symptoms: fever, nasal congestion with/without rhinorrhoea, cough, sore throat, and lymphadenopathy. Nasopharyngeal swabs (NPS) were collected and partitioned for PCR and mNGS assays as described earlier ^5^. In addition to the ARI samples, we collected NPS from four asymptomatic controls (AC). Due to limited sample availability, we excluded one ARI sample from the mNGS assays; this resulted in a total of 53 samples (49 ARI and 4 AC) processed and analyzed in the current study. The median age of ARI participants was 31.9 years and 73.5% of them were female; the median age of AC participants was 35.5 years and 50% of them were female.

After nucleic acid extraction, multiplex real-time PCR was performed using the Amplisens ARVI-screen kit (Amplisens, Moscow, Russia) targeting respiratory RNA (respiratory syncytial virus (RSV), metapneumovirus (MPV), human parainfluenza virus-1-4 (HPIV), coronaviruses (HCoV) OC43/HKU-1 and NL-63/229E and rhinovirus (HRV), and DNA (adenovirus (Adv) B, C and E and bocavirus (BoV) viruses. In 32/49 ARI participants, PCR was positive for HPIV-3 (1/49), HPIV-4 (24/49), HRV (2/49), HPIV4-HRV (4/49) and HAdV (1/49) and HCoV (1/49).

### mNGS

Prior to DNA/RNA extraction and the sequencing library preparation, we enriched NPS samples using a combination of low-speed centrifugation and filtration as previously described ^5^. Free nucleic acids were removed using Pierce Universal Nuclease (Thermo Fisher Scientific, USA). Samples were concentrated using a Pierce ™ PES protein concentrator (Thermo Fisher Scientific, USA) at the 100K molecular weight cutoff. Sequencing library preparation was done as described earlier ^5^. Briefly, nucleic acids were extracted and underwent cDNA synthesis, amplification, and primer removal using the SeqPlex RNA Amplification Kit (Sigma) yielding 150 to 400 nucleotide cDNA fragments, subsequently checked using the NanoDrop and Qubit protocols. Libraries were prepared using the Ion Plus Fragment Library Kit, Ion Xpress Barcode Adapters 1–96 Kit (both from Thermo Fisher Scientific, USA), and the Agencourt AMPure XP kit (Beckman Coulter). Barcoded libraries were assembled using the Ion Plus Fragment Library Kit and Ion Xpress Barcode Adapters 1–96 Kit, and quantified according to the Ion Library TaqMan™ Quantitation Kit’s protocol. Libraries were then loaded onto Ion 530 Chips with the Ion Chef Instrument, and processed on the Ion Torrent S5 System (Thermo Fisher Scientific, USA).

### Bioinformatic analyses

We uploaded the raw fastq files to the Chan Zuckerberg ID (CZID) portal, an open-source metagenomics platform (https://czid.org). The data were standardized based on unique reads mapped per million input reads at the genus tier. To eliminate background and low-frequency sequencing reads, we followed guidelines from the recent metagenomic studies ^13,14^ and set taxon inclusion criteria as follows: nucleotide reads per million (NT %id) of 95% or higher, the alignment length (NT L) of >= 70 base pairs, and reads per million (rpM) >1. We used the NCBI taxonomy browser (https://www.ncbi.nlm.nih.gov/Taxonomy/Browser/wwwtax.cgi) to classify phages according to their bacterial hosts.

### Statistical analyses

Taxon abundance heatmaps were constructed using log-transformed normalized rpM and graphed using Morpheus (https://software.broadinstitute.org/morpheus/). Biodiversity indices were calculated on the normalized rpM data via BiodiversityR, a graphical user interface based on the vegan R package (https://github.com/cran/BiodiversityR), using the default recommended settings for each index. Differences across the PCR and AC sub-groups were assessed using the Mann-Whitney U and Kruskal-Wallis tests in JASP 0.17.2.1 (https://jasp-stats.org/).

### Ethics approval and consent to participate

All study procedures were approved by the Commission on bioethics of KazNARU (dated February, 2023). Written informed consent was obtained from all participants.

## Results

### Nasopharyngeal virome characteristics

mNGS was performed on all (n=53) available samples, with a median of 406,954 reads per sample. After bioinformatic processing, the viral taxa were stratified into human, non-human eukaryotic host, and bacteriophage sub-groups (Fig 1), consisting of a total of 15, 11 and 28 genera, respectively (S1 and S2 Tables). Among the human viruses, gammapapillomavirus and enterovirus were most prevalent in symptomatic participants and observed in 13/49 (26.5%) and 8/49 (16.3%) participants, respectively. No HPIV-4, the virus most prevalently detected by PCR, was detectable by mNGS. Non-human eukaryotic host viruses consisted predominantly of plant and fungal viruses, of which Tobamovirus, associated with tobacco and other Solanaceae, was most prevalent among all participants 23/53 (43.4%). Among the bacteriophages, Pahexavirus, a *Propionibacterium (Cutibacterium) acnes* bacteriophage, was most prevalent and abundant, observed in 48/53 (90.6%) participants and in both ARI and AC groups.

**Fig 1.**
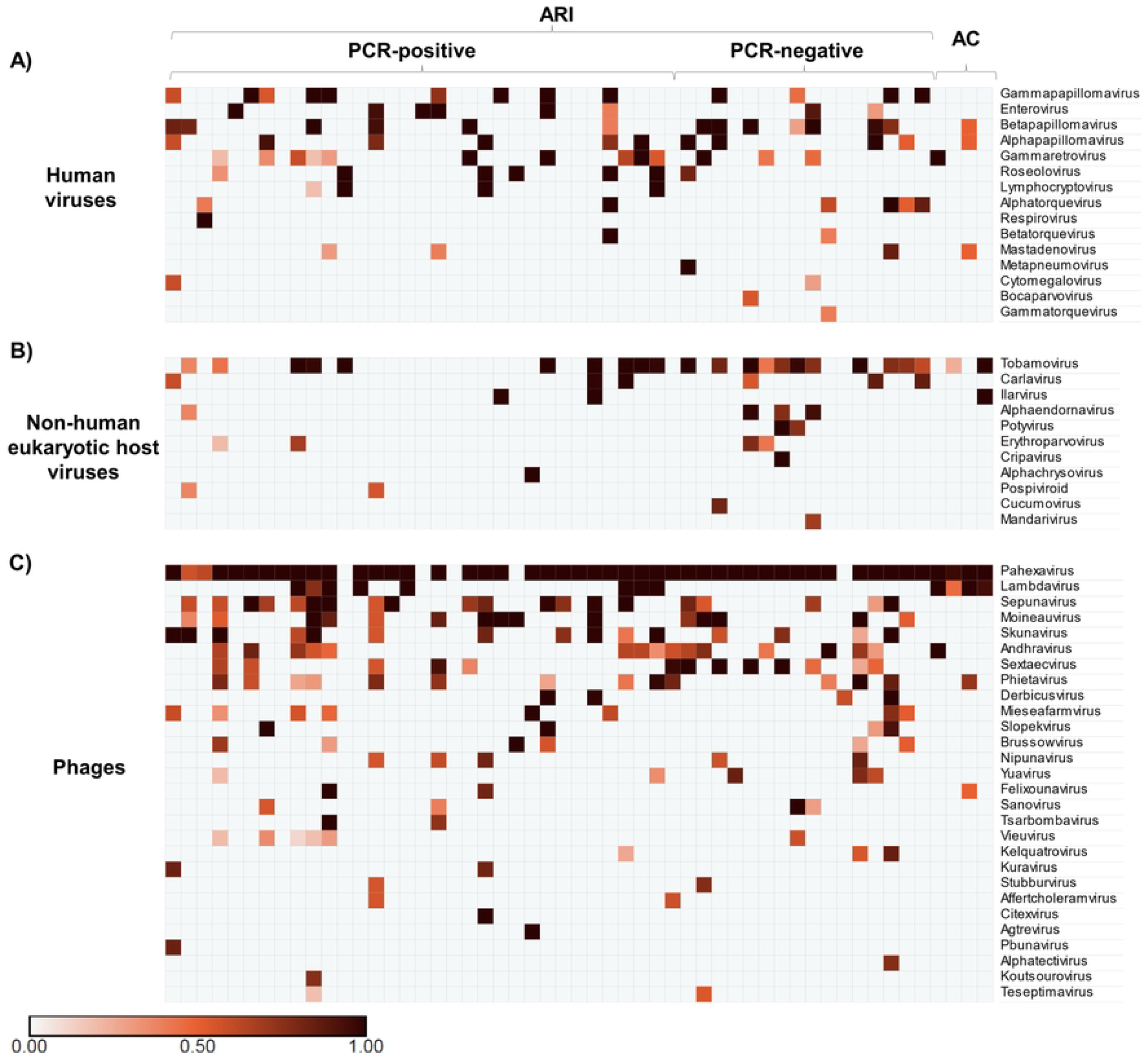
Nasopharyngeal virome profile of participants with acute respiratory symptoms (ARI), stratified by PCR result, and asymptomatic controls (AC). Each row is a viral genus. Each column represents a sample with cells denoting microorganism abundance in reads per million (rpM). A. Human viruses. B. Non-human eukaryotic viruses. C. Phages.

Further, we hypothesized that the nasopharyngeal virome diversity would differ between the participant sub-groups stratified by the multiplex PCR test result. To test this, we calculated two alpha-diversity indices, Simpson’s and Shannon’s, and species richness and compared these indices across the PCR-positive, PCR-negative and AC groups (Fig 2). No significant differences were observed in any of the comparisons (Fig 2).

**Fig 2.**
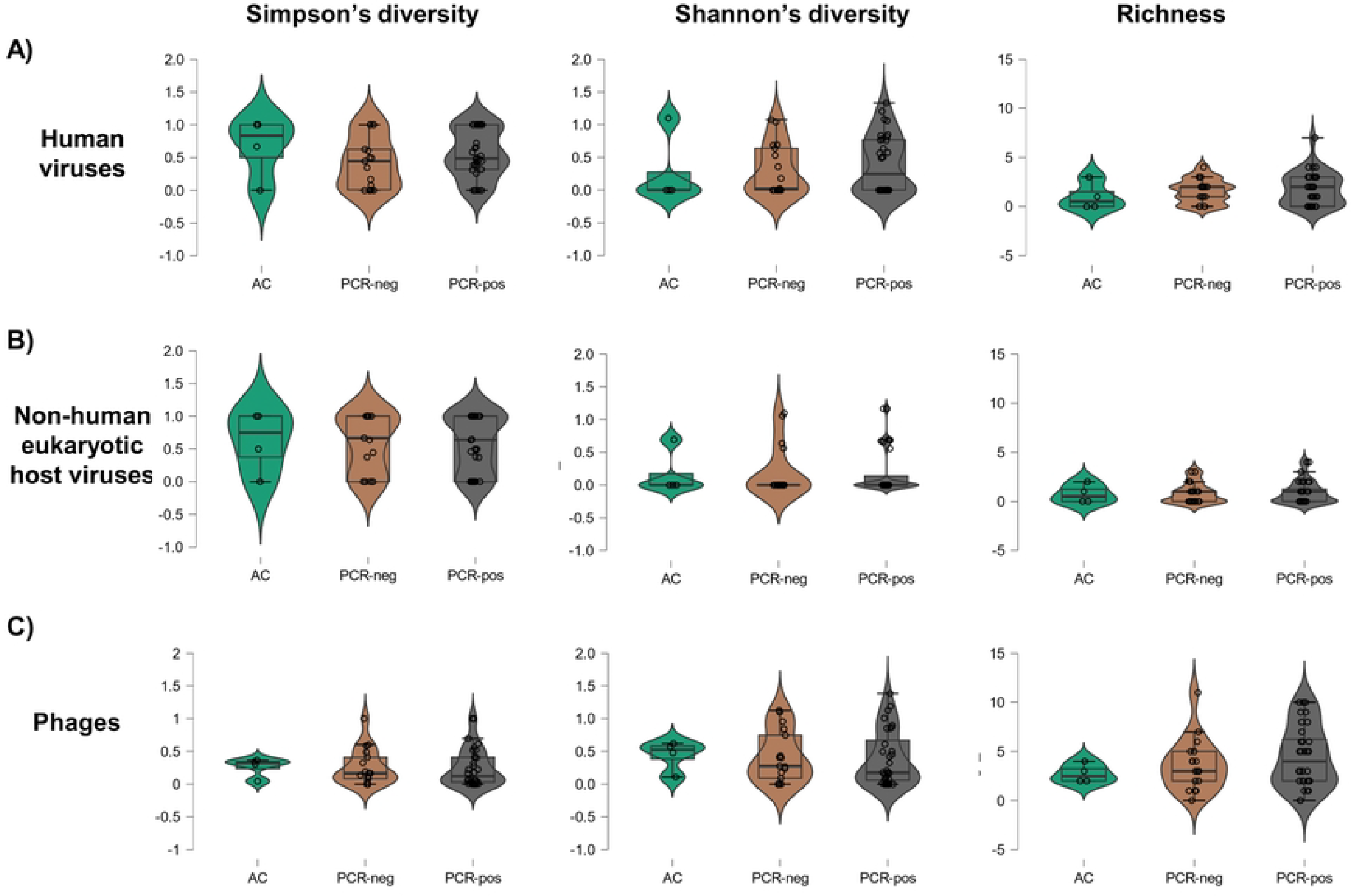
Biodiversity indices for each of the three major viral sub-groups. A. Human viruses. B. Non-human eukaryotic viruses. C. Phages. AC: asymptomatic controls.

### Distribution of ARI-linked viruses and bacteria

We then focused on pathogens known for their associations with respiratory disease, narrowing our analysis to 8 human virus genera and 7 bacterial taxa representing hosts of the 16 recovered bacteriophage genera (Fig 3). Here, we observed a co-detection of viral and bacterial pathogens in 16/49 (32.7%) ARI cases. Only a bacterial or viral pathogen was present in 23/49 (46.9%) and 4/49 (8.2%) ARI cases, respectively. No viral or bacterial pathogens were seen in 6/49 (12.2%) ARI cases (Fig 3). The top three ARI-associated human viruses were enterovirus (16.3%, human rhinovirus, HRV-A), roseolovirus (14.3%, human betaherpesvirus 7, HBV-7) and lymphocryptovirus (8.16%, Epstein-Barr virus, EBV) (Fig 3). The top three ARI-associated phage hosts were *Streptococcus* spp (32.7%), *Pseudomonas aeruginosa* (24.5%) and *Burkholderia* spp. (20.4%). The virome of both asymptomatic and symptomatic subjects was also abundant in the *Staphylococcus* (60.4%) bacteriophages. Except for mastadenovirus and *Staphylococcus*, all other respiratory pathogens were observed only in ARI but not in AC. HRV was detected by mNGS in all PCR+ HRV cases, in addition to two PCR-negative cases and one subject with a PCR-identified HPIV-4 infection. HBV-7 was detected as a viral mono-infection in two cases, and as a co-infection in 5 cases with HRV (n=1), EBV (n=3) and MPV (n=1). EBV was detected in a total of 4 samples, of which only one was a viral monoinfection.

**Fig 3.**
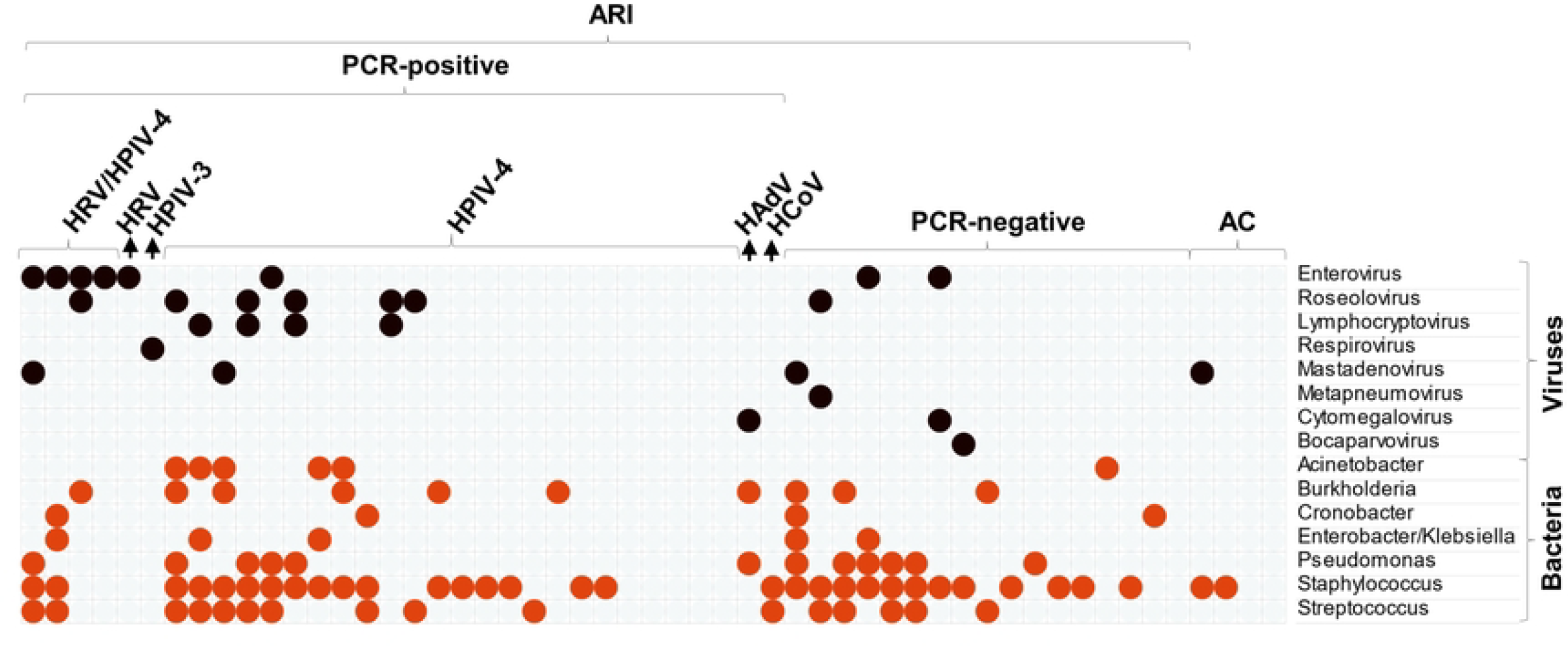
Nasopharyngeal profile of human viruses and bacteria known to cause respiratory disease. Each row is a viral (brown) or bacterial (orange) genus. Each column represents a sample with cells showing presence (colour) or absence of microorganisms. PCR-positive samples are further stratified by the PCR-identified virus. Bacterial genera were derived from the list of mNGS-recovered bacteriophages using phage-host classification. ARI: acute respiratory infection. AC: asymptomatic controls. HRV: human rhinovirus. HPIV: human parainfluenza. HAdV: human adenovirus. HCoV: human coronaviruses (seasonal).

### Concordance between mNGS and PCR

The PCR and mNGS results were relatively concordant for human rhinovirus (HRV) with PCR sensitivity and specificity relative to mNGS of 62.5 (95% CI 24.5, 91.5) and 100% (95% CI 92.1, 100), respectively. The concordance between mNGS and PCR for other PCR panel targets, including human parainfluenza (HPIV), adenovirus (HAdV), bocavirus (BoV) and seasonal coronavirus (HCoV) was low (S3 Table).

## Discussion

In the current study we characterized the nasopharyngeal virome of adults with ARI using mNGS and assess its concordance with a common multiplex PCR assay. In the absence of influenza and respiratory syncytial virus circulation during the study period, our findings were consistent with other studies, where rhinovirus was a leading cause of viral ARI, while *Streptococcus* spp. was the most prevalent ARI-associated bacterial genus ^1^. The identification of roseolovirus and lymphocryptovirus as top ARI-associated human viruses suggests a potential overgrowth of these chronic viruses in the light an ongoing acute infection and mucosal inflammation in adults. Moreover, since HBV-7 is known to primarily cause disease in young children, typically manifesting as a sudden high fever and a distinctive skin rash known as roseola infantum or sixth disease ^15^, our data suggest that adults with ARI can serve as potent HBV-7 reservoirs and transmitters within communities.

Our mNGS data demonstrated a large fraction of the ARI (47%) to be associated with only respiratory bacteria, but not viruses. Notably, half of the mNGS-identified “bacterial-only” samples were also HPIV-4 positive by PCR, suggesting that bacterial load may be overwhelming the molecular signal of viral infection. Cumulatively, these findings support the notion that in clinical practice PCR and mNGS should be used as complementary approaches followed by bacterial culture-based assays. Interestingly, 33% of the ARI in our study were associated with co-detected viruses and bacteria, while viral monoinfections contributed to a small fraction (8%) of the ARI. These findings agree with observations of negative virus-to-virus interactions and positive interactions between viral and bacterial pathogens ^1,16–18^.

To the best of our knowledge, the almost ubiquitous presence in NPS samples of Pahexivirus, a *Propionibacterium acnes* bacteriophage, has not been previously reported. However, a recent 9 thesis ^19^ described a lower relative nasopharyngeal abundance of both Pahexivirus and *Propionibacterium acnes* in HIV+ subjects compared to HIV-negative controls from Malawi, which may reflect an association of *Propionibacterium acnes* with the systemic immune state. We observed a relatively high concordance between PCR and mNGS for HRV but low concordance for other PCR targets, such as HPIV and HAdV. These discrepancies may be attributed to differences between mNGS and PCR in sensitivity, specificity, or the presence of viral variants not detectable by the commercial PCR primers. The absence of HPIV-4 in mNGS, despite detection by PCR, echoes challenges faced in other studies and highlights potential methodological differences or limitations ^8^.

Our study has several limitations. First, the relatively small sample size affects the generalizability of the findings. The low concordance between mNGS and PCR for certain viral targets, such as HPIV and HAdV, highlights potential methodological challenges, possibly related to sensitivity, specificity, or viral variants-which we were unable to investigate within the scope of this work. The large fraction of ARIs associated only with respiratory bacteria, particularly in HPIV-4 positive PCR samples, suggests that further optimization of our mNGS methodology is warranted prior to deployment in clinical settings. We also could not confirm the identified bacterial species as bacterial culture-based assays were not available during the study. In summary, this study sheds light on the multifaceted nature of the nasopharyngeal virome and its potential implications in respiratory health. The discrepancies between mNGS and multiplex PCR results and their clinical significance highlight the need for continued refinement of both techniques and the integration of complementary methods to enhance detection accuracy of ARI pathogens. This work also adds to the growing body of knowledge on the epidemiology of infectious diseases in the region ^20–22^.

## Acknowledgements

We thank all the study participants and the clinic staff.

